# Computational design of monomeric Fc variants with distinct pH-responsive FcRn-binding profiles

**DOI:** 10.1101/2025.05.26.656075

**Authors:** Joon-Young Jeon, Hyejin Lee, Sanghyeon Choi, Seonghyuk Suh, Seong-Bin Im, Yeon-Gil Kim, Bo Mi Ku, Myung-Ju Ahn, Bo-Seong Jeong, Byung-Ha Oh

**Affiliations:** Graduate School of Engineering Biology, Korea Advanced Institute of Science and Technology, Daejeon, Republic of Korea; Therazyne, Inc., Daejeon, Republic of Korea; Department of Biological Sciences, Korea Advanced Institute of Science and Technology, Daejeon, Republic of Korea; Beamline Science, Pohang Accelerator Laboratory, Pohang, Kyungbuk, Republic of Korea; Division of Haematology-Oncology, Department of Medicine, Samsung Medical Center, Sungkyunkwan University School of Medicine, Seoul, Republic of Korea

**Keywords:** Computational design, Monomeric Fc, FcRn, pH-responsive binding affinity, serum half-life extension

## Abstract

IgG1 and IgG4 antibodies form a ∼150 kDa homodimer through dimerization of the Fc domain, which prolongs their *in vivo* half-life via pH-dependent binding to the neonatal Fc receptor (FcRn). Conformationally stable, half-life-extended monomeric Fc (mFc) variants offer a promising platform for antibodies and Fc-fusion therapeutics, enabling deeper tissue penetration, reduced toxicity, and simplified manufacturing. Due to the loss of binding avidity, mFc requires significantly enhanced FcRn-binding affinity at pH 6.0, but retaining weak binding at neutral pH to achieve comparable serum half-life, making engineering such mFc variants highly challenging. Mainly by computational design approach, we created mFc mutants with diverse human FcRn-binding profiles, including two variants that exhibit 17- and 47-fold stronger FcRn binding at pH 6.0 (*K*_D_ of 30.7 nM and 88 nM) compared to a baseline mFc, while maintaining greater than 213-fold weaker binding at pH 7.4 (*K*_D_ of 6,549 nM and 39,150 nM). These variants are highly soluble and display a melting temperature greater than 60.6 °C, underscoring their potential as platforms for extending the *in vivo* half-life of therapeutic modalities. Other mFc variants with different pH-responsive FcRn-binding profiles would potentially fit for other therapeutic needs. Moreover, transferring the same variations into IgG4 Fc generated IgG4 mFc variants with FcRn-binding properties similar to those of the parent IgG1 mFc variants. Furthermore, incorporating the FcRn-binding affinity-enhancing substitutions into native Fc produced a dimeric Fc variant that exhibits strong, pH-responsive FcRn-binding affinities, promising an extended half-life in serum.

## Introduction

Monoclonal IgG1 and IgG4 antibodies are key therapeutic agents. They are a homodimer composed of two heterodimeric subunits, each containing one heavy and one light chain. The fragment crystallizable (Fc) domain of these antibodies consists of two constant heavy chain regions (C_H2_ and C_H3_) and stabilizes the antibody homodimer through C_H3_-C_H3_ interactions. The Fc domain is essential for mediating effector functions, such as prolonging the half-life of IgG antibodies via its interaction with the neonatal Fc receptor (FcRn). FcRn is expressed in various cell types, including endothelial and epithelial cells, and undergoes continuous recycling between the cell surface and endosomes. This recycling is mediated by specific sorting motifs that direct FcRn to recycling or transcytosis pathways, preventing lysosomal degradation [1, 2]. Predominantly localized within the endosomes, FcRn binds endocytosed IgG at low pH (below 6.5) and sorts it away from lysosomal degradation into the recycling pathway [3]. The Fc domain interacts with FcRn in a pH-dependent manner, exhibiting very low affinity at neutral pH (*K*_D_ of ∼88,000 nM) and much higher affinity at low pH (*K*_D_ of 1,377 nM) [4], ensuring rapid release of IgG at the cell surface. A number of studies have shown that the FcRn-binding affinity of Fcs or Fc-containing therapeutic proteins at pH 6 is directly correlated with their serum half-life [5, 6]. Furthermore, the extended half-life of an antibody was shown to correlate with improved *in vivo* therapeutic activity [6, 7].

With an aim to extend the half-life of therapeutic antibodies or Fc-fused therapeutic proteins, Fc variants with enhanced FcRn binding affinity have been discovered by different approaches, including construction and screening of libraries of Fc variants with amino acid substitutions at or near the FcRn-binding interface [8–10]. These include IgG4 Fc variants containing M252Y/S254T/T256E substitutions (YTE) [11], IgG4 Fc variants containing M428L/N434S substitutions (LS) [6] or Q311R/M428L mutations [12]. Interestingly, some of the selected residues do not directly enhance intermolecular Fc-FcRn interactions. For example, in the YTE variant, a prototypic Fc with up to a 10-fold increase in FcRn-binding affinity, the three substitutions were suggested to disrupt intramolecular hydrogen bonds within the C_H2_/C_H3_ domain, increasing side-chain adaptability in residues that form intermolecular contacts with FcRn [13]. Currently, multiple monoclonal antibodies with half-life-extended dimeric Fc variants or therapeutic proteins fused to such Fc are in clinical use [7, 14, 15].

Several studies demonstrated that high-affinity FcRn binding at neutral pH offsets the benefits of increased binding at pH 6.0, likely due to diminished release of antibody into the serum [8, 16, 17]. Therefore, optimizing affinities at pH 6 and pH 7.4 is critical when engineering variants for enhanced serum half-life. Attaining this balance is particularly challenging, as improvements in low-pH affinity often coincide with increases at neutral pH [17]. One study estimated that for a dimeric Fc variant, a neutral pH affinity threshold of approximately 860 nM (*K*_D_) is required to achieve a serum half-life comparable to that of native Fc [8]. Additionally, for dimeric Fc, very low FcRn-binding avidity at neutral pH has been reported as a key factor for longer half-life and achieving optimal pharmacokinetic properties [18].

Generating monomeric Fc (mFc) has garnered significant interest in the biopharmaceutical industry due to its several advantages for clinical use. These include improved tissue penetration, a reduced risk of effector function-related side effects, versatility in creating fusion proteins and its potential use as Fc-only therapeutics to reduce circulating IgG autoantibodies without Fc-mediated effector functions in autoimmune and inflammatory diseases [19]. A number of soluble mFc variants have been developed through approaches such as searching phage-displayed libraries constructed with mutations in the C_H3_-C_H3_ interface [20, 21], applying rational structure-based mutagenesis to weaken the hydrophobic C_H3_-C_H3_ interactions [22, 23] or introducing glycosylation sites into the C_H3_-C_H3_ interface [24].

An mFc itself suffers from a significant loss of FcRn binding activity, as the avidity intrinsic to the bivalent Fc greatly contributes to the strength of FcRn binding [22, 24]. A reported low-pH FcRn-binding avidity of native dimeric Fc is around 300 nM [23], while its low-pH monovalent binding affinity is >2,100 nM [8, 16]. To effectively compete with endogenous IgG for FcRn binding in endosomes, an mFc should exhibit a low-pH binding affinity significantly tighter than 300 nM. While much higher binding affinity at pH 6.0 has been achieved for dimeric Fc, these variants also show enhanced binding at pH 7.4, which can negatively affect serum half-life. Therefore, simply transferring the amino acid substitutions from these dimeric Fc variants to an mFc format would be unsuitable. Unlike their dimeric counterparts, no mFc has yet been adopted for half-life extension in clinically approved therapeutic modalities.

To our knowledge, an IgG4 mFc, named MFc4, maintains *in vivo* terminal elimination half-life of a monomeric bispecific molecule comparable to that of native Fc in human FcRn transgenic mouse, although its serum concentration was lower compared to control IgG. It contains substitutions for disrupting the homodimer interface (L351F/S354E/T366R/P395K/F405R/Y407E) along with additional mutations that are necessary for enhancing FcRn-binding affinity (L428M/L432C/H433S/N434Y/Y436L/T437C/ΔQ438) [21, 23].

We report disruption of the Fc dimer into a conformationally stable mFc based on introducing geometric incompatibility that interferes with the dimeric association of C_H3_ domains. This mFc served as a baseline for computational protein design to enhance the human FcRn-binding affinity in a pH-dependent manner, leading to the creation of seven novel mFc variants. Their FcRn-binding affinities at pH 6.0 vary from 7.5 nM to 145 nM, while at pH 7.4, they vary between 910 nM and 39,150 nM. We further demonstrate that IgG4 mFcs can be generated by incorporating the same amino acid substitutions. Additionally, when a set of FcRn-binding affinity-enhancing substitutions was introduced into the native dimeric Fc, the resulting Fc exhibited significantly increased FcRn-binding avidity at pH 6.0, while retaining weak avidity at neutral pH, a property favorable for extending *in vivo* half-life with good pharmacokinetic properties.

## Results

### Computational design of monomeric IgG1 Fc and experimental validation

The C_H3_ domains of human IgG1 and IgG4 Fc regions contain two short α-helices, α1 and α2. The α2 helix (residues 354–360; SRDELTK) forms a one-and-a-half-turn helix. It is not a standard α-helix, having a tapering C-terminal end. It is somewhat conformationally flexible in both IgG1 Fc and IgG4 Fc, and undergoes a slight but noticeable positional shift upon C_H3_ domain homo-dimerization to avoid steric clashes (Figure 1A). The structure of human IgG1 Fc (PDB entry: 1L6X) [25] showed that incorporating a bulky residue at the Ser354 position on the α2 helix of one subunit would cause severe steric clashes with the other subunit (Figure 1B). We hypothesized that a bulky residue at the Ser354 position might prevent C_H3_ domain dimerization, provided the α2 helix remains rigidly anchored. Using the deep learning-based Inpainting model [26], we generated backbone conformations of α2 with 1 to 4 amino acid insertion(s). Visual inspection indicated that a single-residue insertion between Pro354 and Gln362 would be optimal, as it resulted in a regularly shaped ɑ-helix without perturbing the loop connection to the following ꞵ-strand (Figure 1C). Subsequently, sequences for the extended α2 helix were designed using the deep learning-based ProteinMPNN model [27]. Fc dimeric structure prediction of these designs by using AlphaFold2 (AF2) [28] showed a close resemblance between the designed α2 model and the predicted models. From these, six designs were selected for experimental validation. In all these sequences, residue 354 was deliberately substituted with arginine to sterically hinder C_H3_ dimerization. Additionally, Tyr407, a key residue involved in intermolecular hydrophobic interactions, was intentionally replaced with glutamate to reduce the hydrophobicity of the dimer interface region, thereby improving solubility [21].

**Figure 1.**
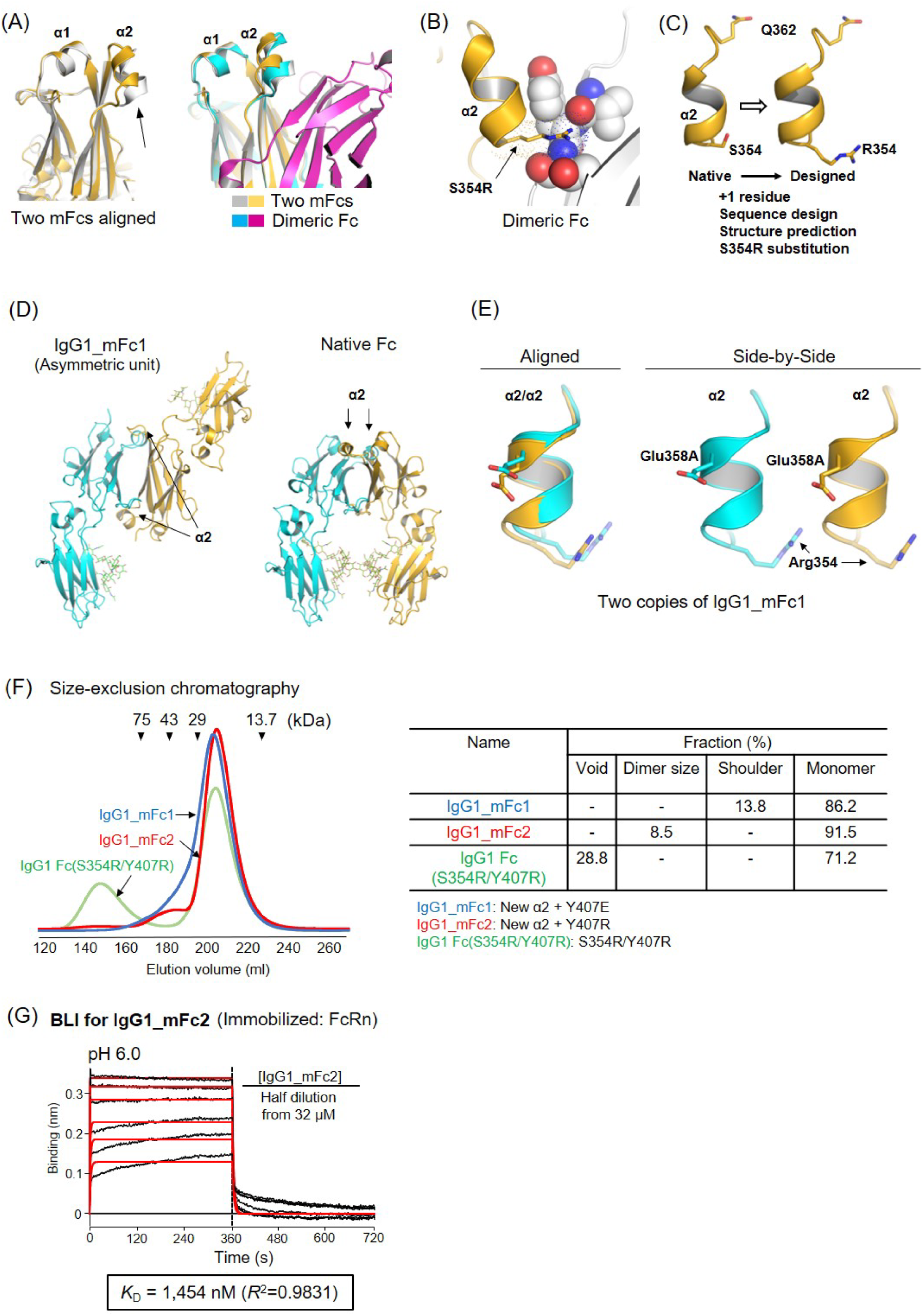
Design and experimental validation of IgG1_mFc1 and IgG1_mFc2. (A) Conformational flexibility of helix α2. Superposed are the C_H3_ domain in two independent IgG1 mFc molecules in the asymmetric unit of a crystal (PDB entry: 6F2Z), highlighting positional and conformational variability of α2 (*Left*). The same mFc molecules superposed on native dimeric IgG1 Fc (1L6X), showing positional adjustment to form the homodimer (*Right*). The α2 sequences are identical in both the monomeric and dimeric structures. (B) Dimer-disruptive mutation. S354R substitution at the beginning of α2 in the IgG1 structure (PDB entry: 1L6X) is not sterically tolerable. The two subunits are shown in surface representation with different colors and Arg354 is shown in stick representation. (C) Remodeling of helix α2. The α2 helix in 1L6X was extended by one residue using Inpainting, followed by sequence design with ProteinMPNN and semi-validation via AF2. For comparison, the structural model of a designed mFc, incorporating the S354R substitution, is shown on the right, displaying a regular α-helix formation. (D) Disruption of native homodimer formation. Two molecules in the asymmetric unit of IgG1_mFc1 are displayed alongside the native Fc homodimer (PDB entry: 1L6X). (E) Conformation of α2 in IgG1_mFc1. The C_H3_ domain in two copies of IgG1_mFc1 molecules in the asymmetric unit were superposed (RMSD: 0.25 Å), and only the two α2 helices are shown together (*Left*) and side-by-side after translation (*Right*), illustrating their closely similar regular helical conformations. Shown in sticks are the strategically introduced Arg354 and the inserted residue Glu358A, which is solvent-exposed. (F) Solution behavior. The indicated Fc variants, after elution from protein A resin, were concentrated to 10 mg/mL and loaded onto a Sephadex 75 column. Size marker proteins are indicated by triangles. (G) BLI run. Binding affinity of IgG1_mFc2 for FcRn was measured at pH 6.0, and the deduced affinity is shown. Fitted curves are displayed in red throughout the figures, and the corresponding *kₐ* and *k_d_* values are provided in Supplementary Figure 1.

The six selected designs were expressed in Chinese hamster ovary (CHO) cells and analyzed by size-exclusion chromatography. The design with the best elution profile (a major single peak with a tailing shoulder) was selected and designated as IgG1_mFc1. The elution peak corresponded to the monomer size of IgG1 Fc. For further confirmation of the monomerization, we determined the crystal structure of IgG1_mFc1 (Supplementary Table 1). In the crystal structure, the homo-dimerization interaction observed in the native Fc is disrupted because the two molecules of IgG1_mFc1 contained in the symmetric unit do not form a dimer (Figure 1D). This observation confirms that IgG1_mFc1 remains monomeric in the crystalline state where the IgG1_mFc1 concentration should be high. The conformation and positioning of α2 in the two IgG1_mFc1 molecules within the asymmetric unit are nearly identical (Figure 1E), indicating that the α2 is stably anchored to the main body of Fc and Arg354 prevents the dimerization interaction via geometric incompatibility, as intended. Structural alignment showed that α2 in IgG1_mFc1 has a single residue insertion and five substitutions compared to the native Fc; 354-<u>SRD</u>ELT<u>KN</u>-361 is replaced by 354-<u>RPE</u>EL**E**T<u>QE</u>-361 with the inserted residue in bold and substituted residues underlined. To maintain consistency with the native sequence numbering, the internal LET residues are numbered as Leu358, Glu358A and Thr359.

We tested whether substituting Y407R for Y407E could improve the solution behavior of IgG1_mFc1. Indeed, this change decreased the tailing shoulder from 13.8% to 8.5%, indicating that Y407R is more effective for Fc monomerization (Figure 1F). The resulting variant was designated IgG1_mFc2. Notably, IgG1 Fc with only the S354R/Y407R substitutions, IgG1 Fc(S354R/Y407R), eluted also as a monomer but showed a significant level of soluble aggregates (Figure 1F). As detailed below, this variant also had a markedly lower melting temperature than IgG1_mFc2, indicating that the remodeled α2 helix contributes considerably to protein folding and stability. Next, we evaluated the FcRn-binding affinity of IgG1_mFc2 via biolayer interferometry (BLI) at pH 6.0 and obtained a *K*_D_ value of 1,454 nM (Figure 1G), which is comparable to the reported 2,140 nM for native dimeric IgG1 Fc [8]. Similar *K*_D_ value (2,930 nM) was obtained for IgG1_mFc1 (Table1; Supplementary Figure 1).

### Computational design to enhance FcRn-binding affinity of IgG1_mFc2

Structural studies suggested that His310 and His435 play a pivotal role in pH-responsive binding to the FcRn receptor [29, 30]. Amino acids near these two residues have been the primary focus to discover Fc variants with enhanced FcRn-binding affinity, preferentially at lower pH, by rational mutagenesis or phage display library screening. Since these efforts have been extensive, we sought an approach distinguished from the previously reported methods. The first target was the His310^Fc^-Glu115^FcRn^ interaction. The crystal structure of Fc(YTE)‒ albumin‒FcRn (PDB entry: 4N0U) [30] shows that the imidazole ring of His310^Fc^, which is surrounded by carbonyl oxygens, appears to make an intermolecular interaction with Glu115^FcRn^ in a pH-responsive manner: tighter ionic interaction at low pH (Asp^⊖^···His^⊕^) than the electrostatic interaction at neutral pH (Asp^⊖^···His). Nearby hydrophobic residues close to this interaction are Ile253 and Leu309 of Fc, Trp328 of the FcRn heavy chain and Ile1 of ꞵ2M (Figure 2A). We postulated that increasing hydrophobicity (thereby decreasing dielectric constant) near the His310^Fc^-Glu115^FcRn^ interaction pair would reinforce their intermolecular interactions preferentially at low pH. We then focused on a loop (residues 284-286) following ꞵ4 and spanning one edge of the C_H3_ domain of Fc, hypothesizing that extending this loop could bring it closer to the His310^Fc^-Glu115^FcRn^ interaction, with newly introduced hydrophobic residues potentially enhancing the hydrophobic environment around the interaction pair. Using RF Diffusion, numerous extended backbones were generated with Ile1 and Gln2 of ꞵ2M designated as hotspot residues. Among these, those forming a half-turn α-helix were selected for sequence design using ProteinMPNN. Nearby residues were also varied during the sequence design process, followed by structure prediction with AF2 (Figure 2B). A single design with high AF confidence scores was chosen for experimental validation by quantifying the FcRn-binding affinity by BLI. In this selected design, designated as IgG1_mFc3, the sequence 284-VHNA-287 of the native IgG1 Fc was replaced with 284-<u>AE</u>**KVGF**<u>VP</u>-287, where KVGF are the inserted residues and the underlined residues are the substituted ones. To maintain consistency with the native sequence numbering, the internal EKVGFV residues are designated as Glu285, Lys285A, Val285B, Gly285C, Phe285D and Val286. Size-exclusion chromatography showed IgG1_mFc3 eluting as a single, monodisperse peak (Figure 2C). Importantly, BLI runs at multiple IgG1_mFc3 concentrations showed a marked increase in FcRn-binding affinity, improving from a *K*_D_ of 1,454 nM of IgG1_mFc2 to 88 nM at pH 6.0. The measured FcRn-binding affinity of this variant at pH 7.4 was 39,150 nM, corresponding to 445-fold difference in the binding affinity between the two pH values (Figure 2D). The FcRn-binding affinity of IgG1_mFc2 at pH 7.4 was too weak to reliably measure in our assay. However, compared to the reported affinity (*K*_D_ of 88,000 nM) for dimeric IgG1 [16], its measured affinity of 39,150 nM represents only 2.2-fold increase. We determined the structure of IgG1_mFc3 (Figure 2E), in which the extended loop forms a half-turn α-helix and is conformationally rigid, as evidenced by well-defined electron densities. Its conformation closely matches the design model, in which it interacts with FcRn and ꞵ2M as well as mFc, explaining the enhanced binding affinity. The huge affinity difference between the two pH values may result from reinforcing the electrostatic interaction between the protonated His310^Fc^ and Glu115^FcRn^ at low pH, as we intended.

**Figure 2.**
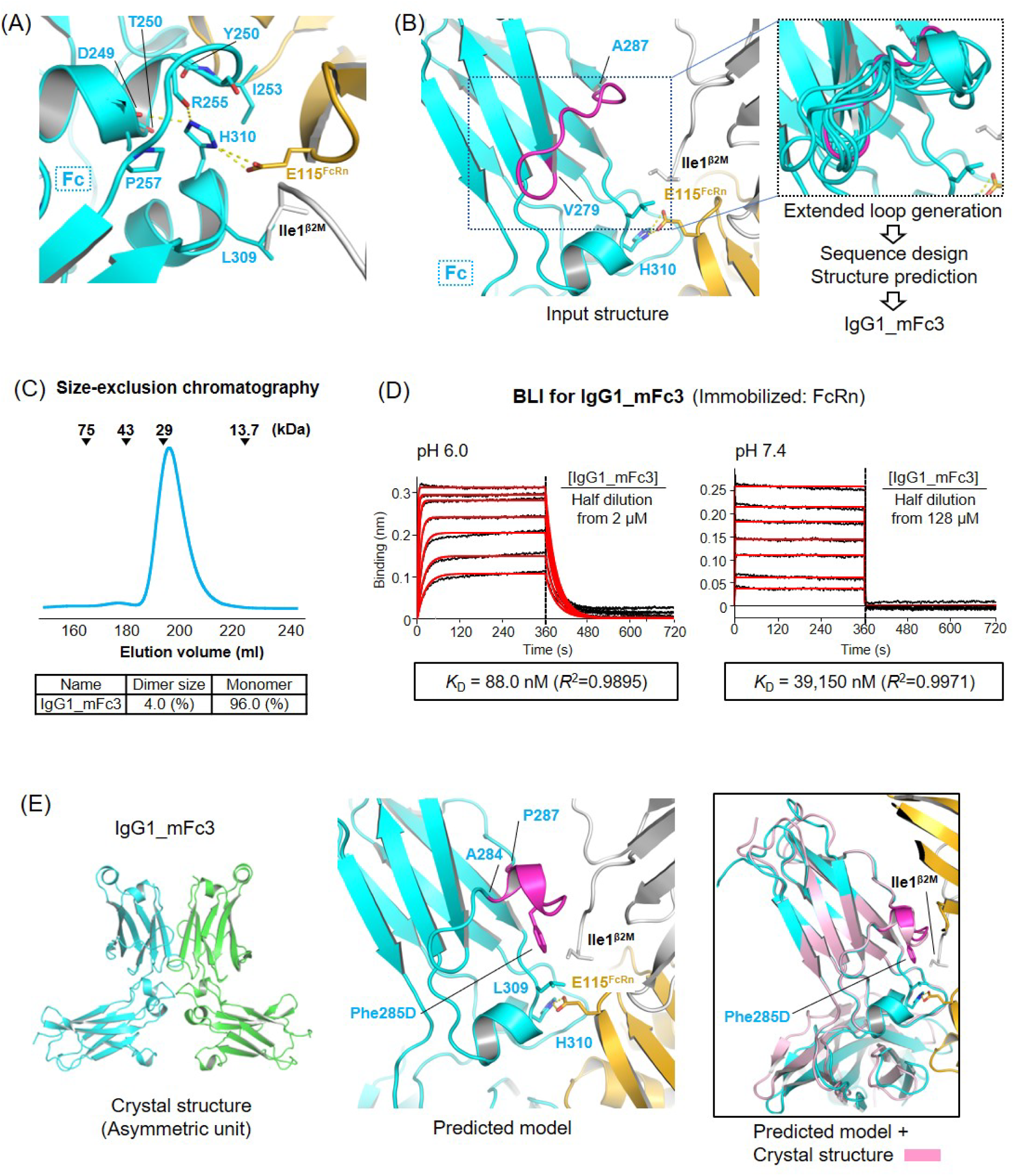
Computational design and characterization of IgG1_mFc3. (A) Environment around the His310^Fc^-Glu115^FcRn^ interaction pair. The ternary complex of IgG1 Fc(YTE)‒albumin‒FcRn is shown. The imidazole ring of His310^Fc^ is positioned near the carbonyl oxygens of Asp249, Thr250, Tyr252, Ile253 and Arg255. Those of Asp249 and Arg255 form hydrogen bonds (yellow dots) with His310. The hydrophobic side chains (Ile253^Fc^, Pro257^Fc^, Leu309^Fc^, Ile1^ꞵ2M^), shown in stick representation and labeled, partially bury the His310^Fc^-Glu115^FcRn^ interaction. (B) Computational procedure. The coordinates of 6WNA served for the loop extension within residues 279-287 of Fc (in magenta) by using RF diffusion, followed by sequence generation by ProteinMPNN and structure prediction by AF2. Based on the AF confidence scores, a single design was selected (IgG1_mFc3). (C) Solution behavior. IgG1_mFc3, after elution from protein A resin, was concentrated to 10 mg/mL and loaded onto a Sephadex 75 column. Size marker positions are shown. (D) Quantification of binding affinities. Binding affinity measurements were performed using BLI. Experiments were conducted in technical triplicates, producing consistent results. Representative sensorgrams and the calculated *K*_D_ values are shown. (E) Crystal structure and binding model. The crystal structure of IgG1_mFc3 is shown (*Left*). The AF2-predicted FcRn binding model shows that Phe285D residue on the extended loop (highlighted in magenta) augments the hydrophobic interaction between Fc’s Leu309 and Ile1^ꞵ2M^ (*Middle*). An overlay of the predicted structure onto one subunit in the crystal structure; the Cα atoms of residues 238–345 were superposed with an RMSD of 0.34 Å, demonstrating that the extended loop portion (in magenta) adopts a similar conformation (*Right*).

### Affinity tuning by targeting the His435 region

We aimed to identify other amino acid substitutions to further enhance the FcRn-binding affinity of IgG1_mFc3, focusing on the region surrounding His435 of Fc, which also plays a role in pH-responsive FcRn binding. Asn434-His435 participates in intermolecular interactions with FcRn; the side chain of Asn434 forms hydrogen bonds with carbonyl oxygens of Gly129^FcRn^ and Trp131^FcRn^, while the imidazole ring of His435, surrounded by the carbonyl oxygens of Leu251^Fc^, Glu430^Fc^, Leu432^Fc^ and Asp130^FcRn^, makes van der Waals contact with Trp131^FcRn^ and Pro132^FcRn^ (Figure 3A). Previous studies showed that M428L/N434S mutations in Fc enhance FcRn-binding affinity [6]. In the case of M428L, substituting leucine at this position appeared to affect the hydrophobic interactions with Leu251^Fc^, Met252^Fc^ and Pro132^FcRn^, thereby indirectly influencing the pH-responsive interaction of His435 (Figure 3A). We examined whether the M428L substitution in IgG1_mFc3 could enhance the FcRn-binding affinity, and found that it indeed improved the affinity from a *K*_D_ of 88 nM to 31 nM at pH 6.0, while also increasing the binding affinity approximately 6-fold at pH 7.4 from a *K*_D_ of 39,150 nM to 6,549 nM (Figure 3B; *Left*). Therefore, the affinity enhancement by the M428L substitution in the IgG1_mFc3 background is based toward neutral pH. We designated this variant as IgG1_mFc4. Next, based on the structure, we hypothesized that N434H substitution could introduce an imidazole ring interaction with Leu135^FcRn^ (Figure 3C), thereby enhancing FcRn-binding affinity further. Binding experiments indeed showed that the additional N434H substitution enhanced the affinity at pH 6.0, reducing the *K*_D_ from 30.7 nM to 7.5 nM (∼4-fold increase), and at pH 7.4, from 6,549 nM to 1,056 nM (∼6-fold increase) compared to IgG1_mFc4 (Figure 3B; *Middle*). We designated this variant as IgG1_mFc5.

**Figure 3.**
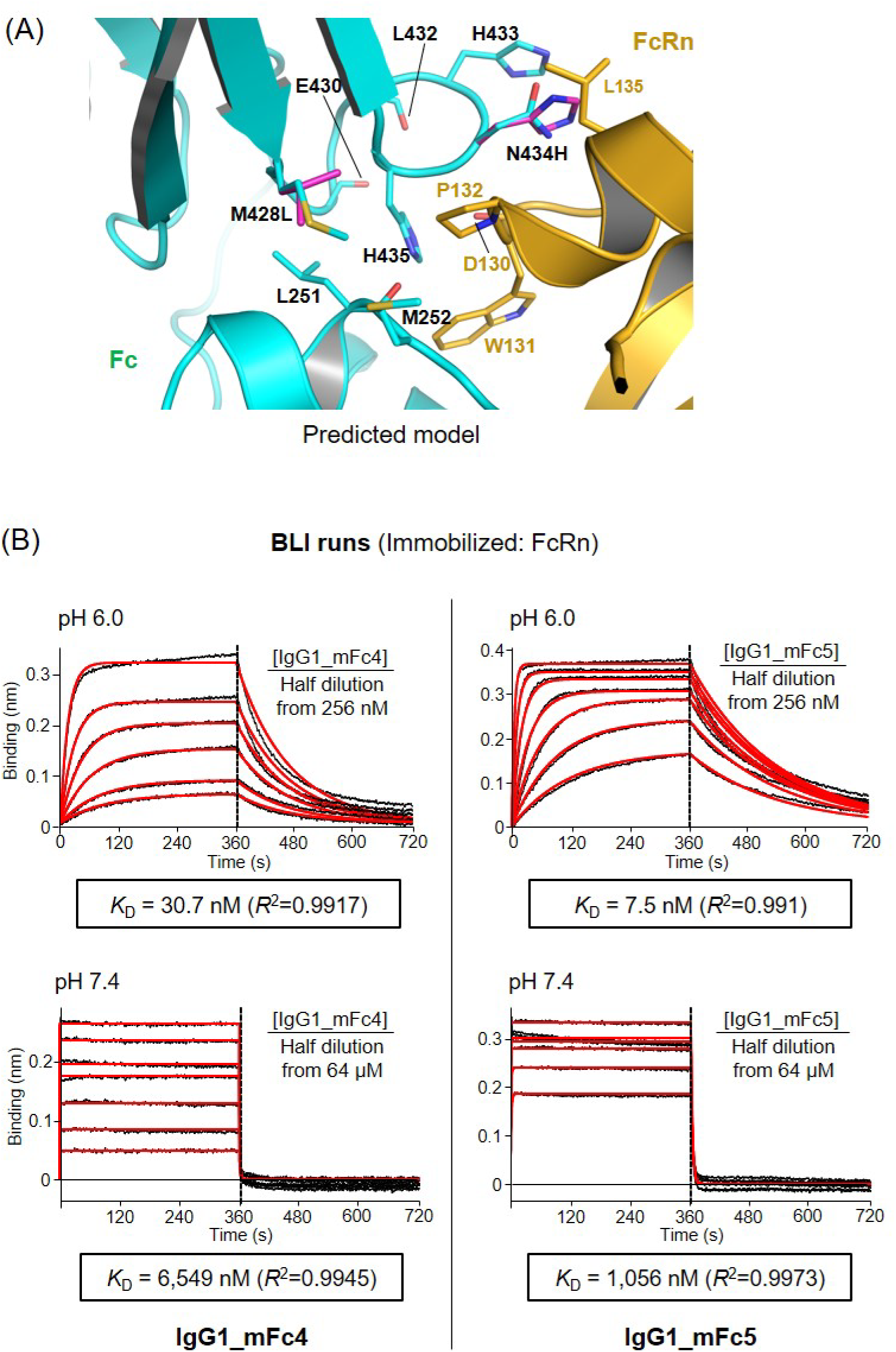
Modification targeting the His433-Asn434-His435 loop of Fc. (A) Environment around His435^Fc^. In the AF2-predicted structural model of IgG1_mFc3 bound to FcRn–ꞵ2M, the environment around His435^Fc^ is highlighted. Four carbonyl oxygens (Leu251^Fc^, Glu430^Fc^, Leu432^Fc^, Asp130^FcRn^) and four hydrophobic residues (Leu251^Fc^, Met428^Fc^, Trp131^FcRn^,Pro132^FcRn^) close to the imidazole ring of His435^Fc^ are shown in sticks and labeled. Two amino-acid substitutions using the PyMOL software are shown in magenta. Note that the imidazole ring of His434^Fc^ stacks with that of His433^Fc^ and lies close to the hydrophobic side chain of Leu135^FcRn^. This imidazole ring stacking between His433^Fc^ and the substituted residue His434^Fc^ is observable in the crystal structure of IgG1_mFc9, described below. (B) BLI runs. The binding affinities of the three indicated mFc variants were measured at pH 6.0 and pH 7.4, and the deduced affinities are shown.

### Further affinity tuning by combining T256Y, T307L and V308I substitutions

Increased conformational dynamics of the C_H2_-C_H3_ interface of Fc have been shown to enhance FcRn-binding affinity [13]. Based on this, we aimed to modulate the FcRn-binding affinities of IgG1_mFc4 and IgG1_mFc5. We selected three residues, Thr256, Thr307 and Val308 in Fc, which are located on the His310-containing segment (residues 307-319) and its interacting loop (residues 252-258) (Figure 4A). T256 and T307 are exposed residues, and the T256Y and T307L substitutions were expected to bring them into contact. V308 is a buried residue, and the V308I substitution, found in rabbit Fc, was anticipated to fill the internal void and enhance the packing of buried apolar residues. Together, these might increase the conformational rigidity of the two segments. The three substitutions were introduced into IgG1_mFc4 and IgG1_mFc5, generating variants IgG1_mFc6 and IgG1_mFc7. We confirmed that the expected local changes by the three substitutions were confirmed by the structure determination of IgG1_mFc6, which contains all three substitutions (Figure 4A; Supplementary Table 1). These additional substitutions mostly decreased the binding affinity at both pH 6.0 and pH 7.4. As a result, IgG1_mFc6 exhibits strong pH-dependency, with a *K*_D_ of 44 nM at pH 6.0 and 11,000 nM at pH 7.4 (Figure 4B; Table 1).

**Figure 4.**
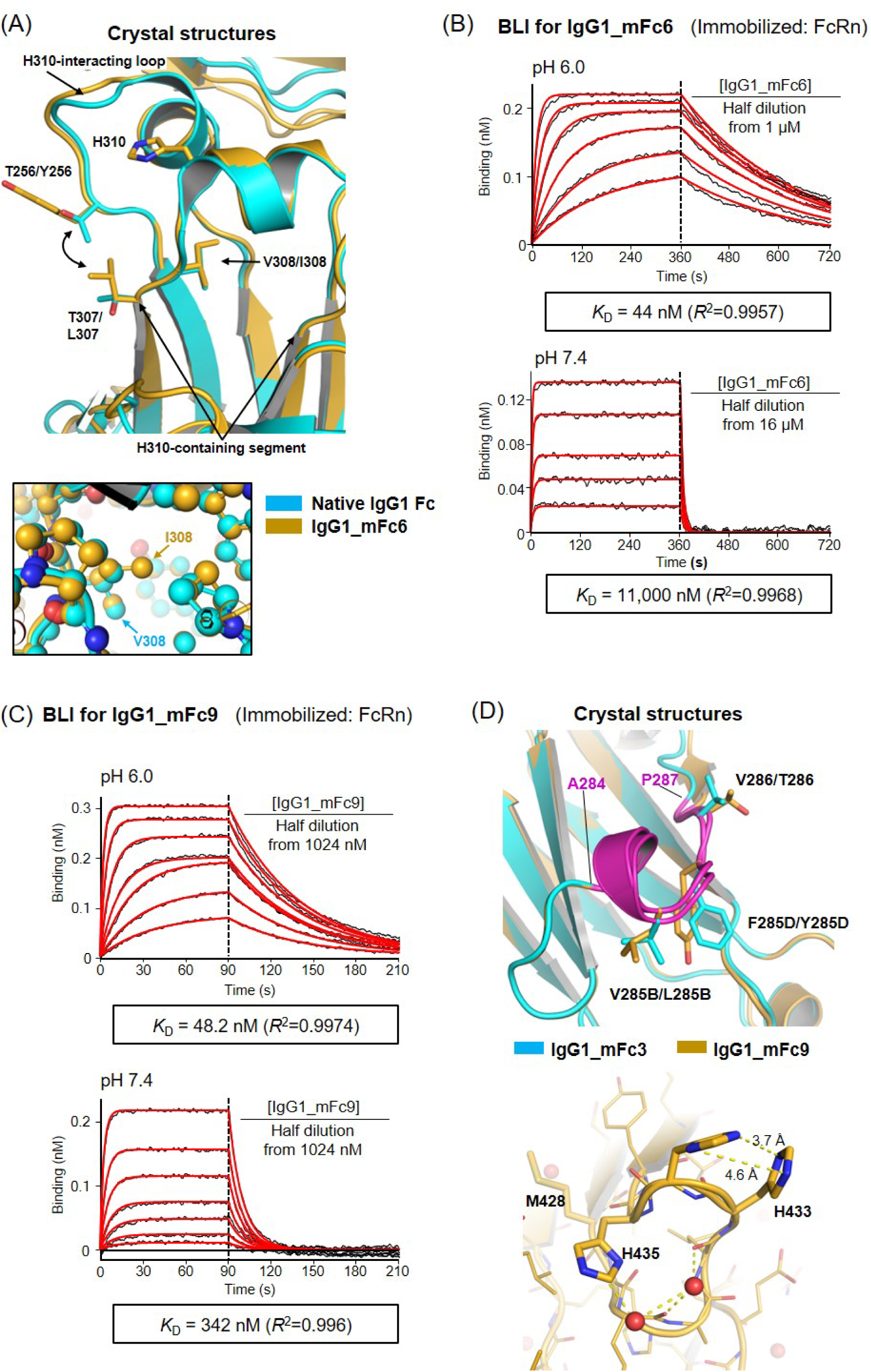
Affinity tuning based on amino acid substitutions near His310 or on the designed loop. (A) Locations of T256, T307, V308. The crystal structure of native IgG1 Fc (PDB entry: 1L6X) was superimposed onto that of IgG1_mFc6. The three varied amino acids and His310 are shown as stick representations. Arrows indicate the start and end of the His310-containing segment. The hydrophobic contact between Y256 and L307 is indicated by the double-headed arrow. The inset shows the void filling by the extra methyl group of I308 in IgG1_mFc6 compared to V308 in the native Fc. The radii of atoms are deliberately reduced. (B) BLI runs for IgG1_mFc6, along with the deduced *K*_D_ values, are shown. (C) BLI runs for IgG1_mFc9, along with the deduced *K*_D_ values, are shown. (D) Structural features and FcRn-binding affinity of IgG1_mFc9. The IgG1_mFc9 structure is superposed onto that of IgG1_mFc3. The designed loop (residues A284-P287) is shown in purple, and three amino acid variations within this loop are highlighted in sticks (*Left*). The H435-containing loop is shown (*Right*). The substituted H434 is involved in an imidazole ring stacking interaction with His433. Distances between ring nitrogen atoms are indicated. Bound water molecules and their hydrogen bonds are represented by spheres and dotted lines, respectively.

**Table 1.**
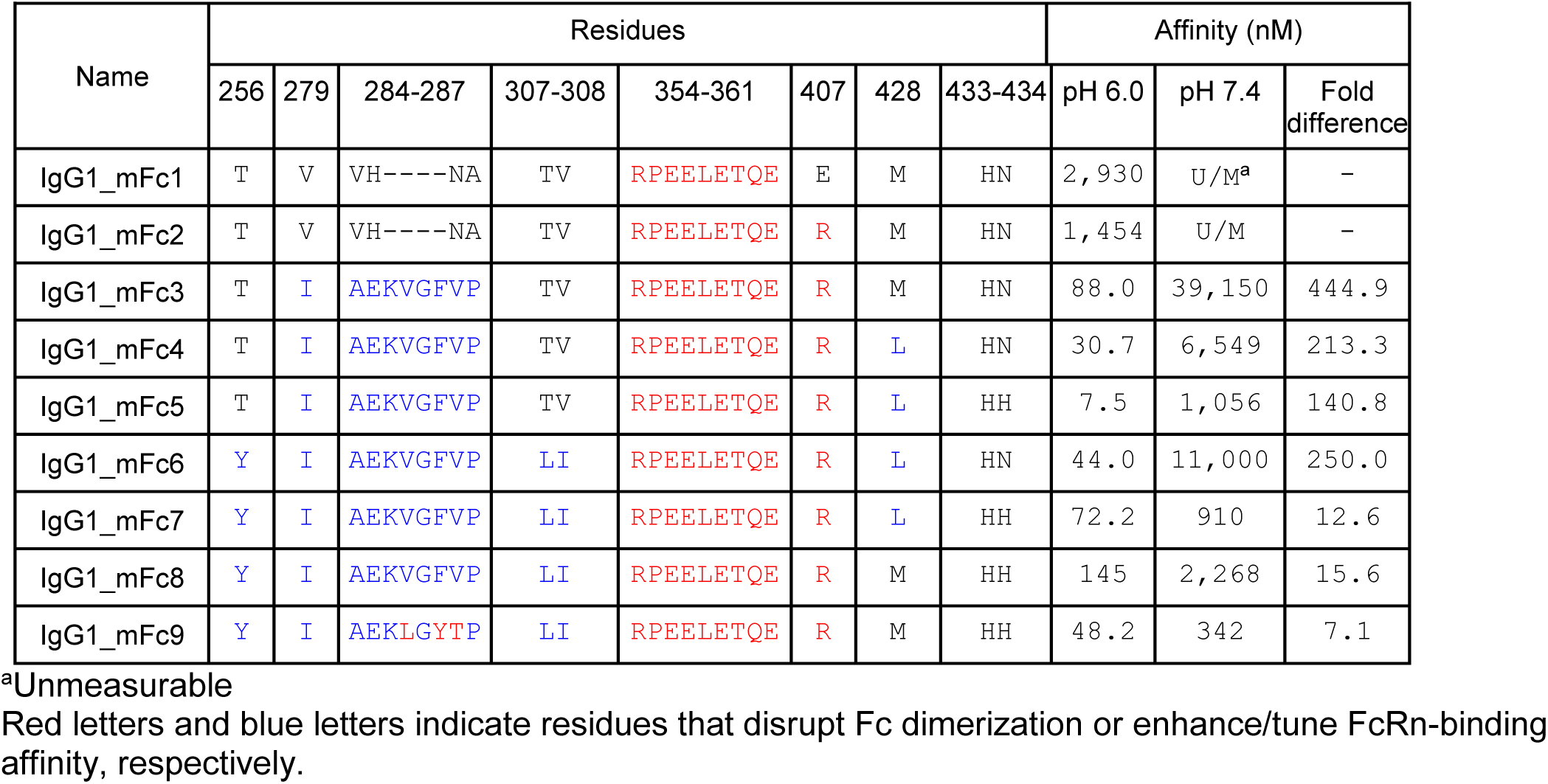
Varied sequences and FcRn-binding affinities of mFc variants generated in this study.

### The designed loop as a source for generating new variants

While it is possible to tune the FcRn-binding affinity without altering the designed loop (284-AEKVGFVP-287), we expect that sequence variation within this loop could serve as a rich source for generating monomeric and dimeric Fc variants with diverse FcRn-binding profiles. Supporting this, the variant IgG1_mFc9, differing from the parent IgG1_mFc8 in three amino acids within the loop (Val285B→Leu, Phe285D→Tyr, Val286→Thr), exhibited markedly different FcRn-binding affinities (*K*_D_ of 48.2 nM and 342 nM) from those of IgG1_mFc8 (*K*_D_ of 145 nM and 2,268 nM) (Figure 4C; Table 1; Supplementary Figure 1). We determined the crystal structure of IgG1_mFc9 (Figure 4D), which shows that the extended loop in this variant is also conformationally rigid. While neither the IgG1_mFc9 structure nor a predicted model of its complex with FcRn–β2M (not shown) explains the differential affinity increase compared to IgG1_mFc8, the amino acid substitutions in the designed loop likely alter the electrostatic environment around the His310^Fc^-Glu115^FcRn^ pair in an unpredictable manner. Screening a random amino acid substitution library of this loop would be a promising approach to discover mFc variants with widely varying pH-responsive FcRn affinities.

### Influence of protein fusion to mFc

To assess whether protein fusion to mFc variant affects the FcRn-binding affinity, we generated IgG1_mFc3 fused to the Fab domain of Trastuzumab, an anti-HER2 IgG1 antibody. The FcRn-binding affinities of this fusion protein (68.7 nM at pH 6.0 and 17,000 nM at pH 7.4) were comparable to those of IgG1_mFc3 alone (88 nM and 39,150 nM) (Supplementary Figure 1, bottom), indicating that the fused Fab domain negligibly affect the mFc interaction with FcRn.

Next, to assess whether mFc fusion affects the production of a client protein, we compared the expression yield of human growth hormone (hGH) alone and hGH fused to the N-terminus of two different mFcs. Although the purification methods are different between the two constructs, the productivity in total amount of hGH-mFc fusion proteins was higher than hGH alone in transient expression of the proteins. However, normalized productivity according to the molecular weight showed that the final yield of the fusion proteins was about 25% less than hGH alone (Supplementary Figure 4). These comparisons indicate that the mFc fusion negligibly affect the expression of the client protein.

### Protein stability and immunogenicity of mFcs

Antibody stability is critical, as it directly impacts efficacy, half-life and safety. We evaluated the thermal stability of four mFcs (IgG1_mFc2 through IgG1_mFc5) by measuring their melting temperature (*T*_m_) using circular dichroism (CD) spectroscopy (Figure 5A). They exhibited melting temperatures of 58.4-62.2 °C, which is higher than or at least comparable to the typical 45–58 °C range reported for most mFcs, unless an additional disulfide bond is introduced [21, 23, 24, 31].

**Figure 5.**
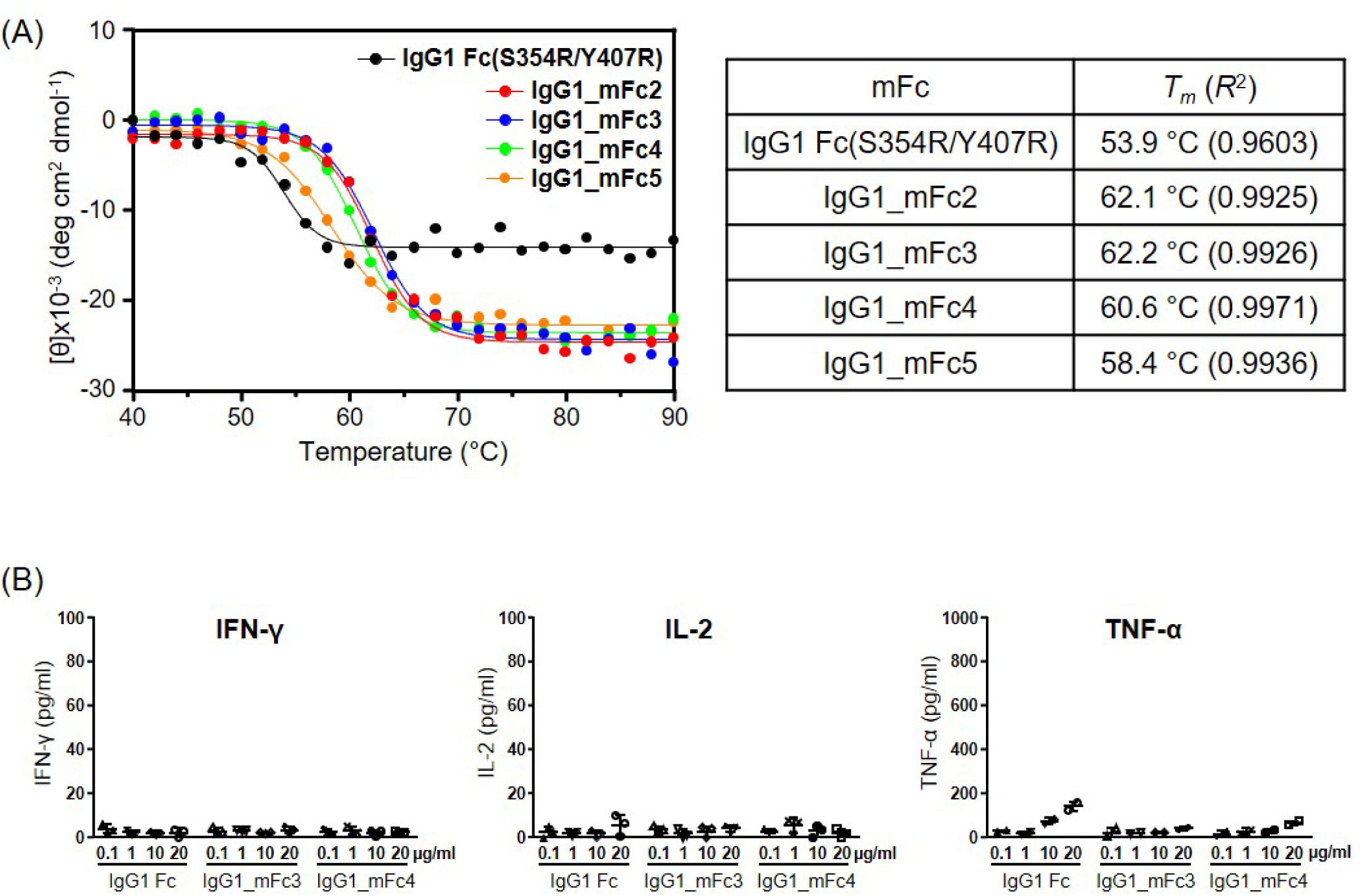
Thermal stability and immunogenicity. (A) Melting temperature (*T*_m_) measurement. The CD ellipticity at 206 nm was measured as a function of temperature for the indicated mFcs. For IgG1 Fc(S354R/Y407R), the monomer-size fractions from the size-exclusion chromatography column were collected and used. The *T*_m_ values and the *R*^2^ values for curve fitting are shown. (B) PBMCs at a concentration of 1×10^6^ cells/mL were stimulated for 24 h with native IgG1 Fc or the three indicated mFc variants. Culture supernatants were collected and analyzed by ELISA to measure IFN-γ, IL-2, and TNF-α levels. The bar graphs represent the mean ± SD from PBMCs derived from two or three individuals. No significant differences in cytokine secretion were observed between the native IgG1 Fc and any of the mFc variants.

Artificial variations on the Fc domain may result in new T cell epitopes that are processed and presented by antigen-presenting cells, leading to T cell-mediated immune responses. We evaluated the immunogenicity of two mFc variants by measuring the secretion of inflammatory cytokines (IFN-γ, IL-2, TNF-α) using enzyme-linked immunosorbent assay (ELISA). The variants were IgG1_mFc3 (with the additional designed loop), and IgG1_mFc4 (with the additional designed loop plus the M428L variation). The two mFcs induced no noticeable secretion of the cytokines similar to the native IgG1 Fc (Figure 5B).

### Comparison with other reported mFc variants

The amino acid variations and substitution positions in our mFc variants are distinct from those reported in the literature (Table 2; shaded area), underscoring the uniqueness of our approach to Fc dimer disruption and the enhancement of FcRn-binding affinity. To our knowledge, MFc4 is the only Fc variant reported to exhibit a single-digit nanomolar FcRn-binding affinity, with a reported *K*_D_ of 5 nM at pH 6.0 [23]. In our mFc variants, the amino acid substitution positions do not overlap with those in MFc4 except at residue 354, where our variants contain S354R compared to S354E in MFc4 (Table 2).

**Table 2.**
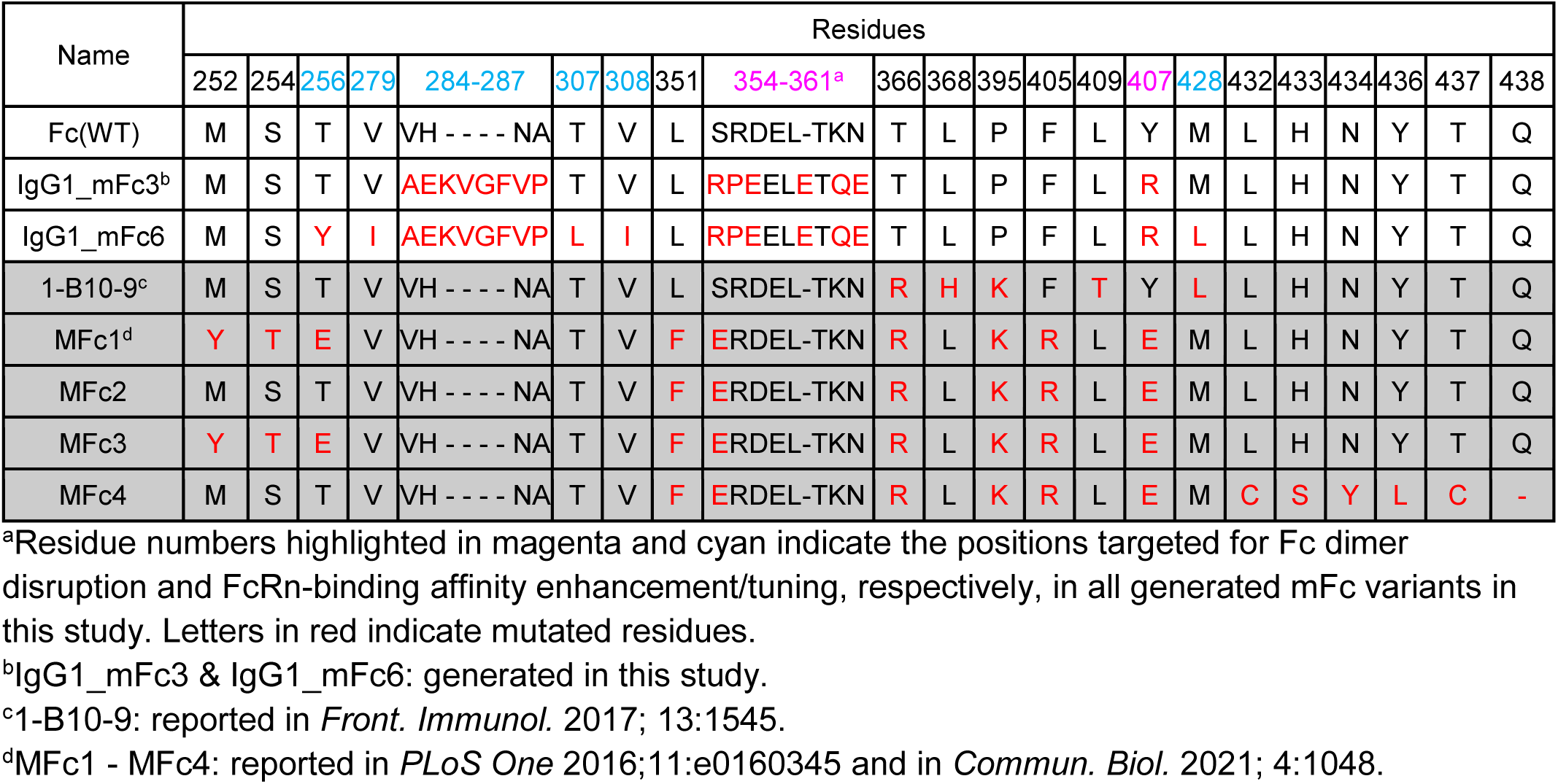
Comparison of the mutational positions in mFc variants.

Since the FcRn-binding affinity of MFc4 at neutral pH was not previously reported, we prepared this variant and measured their FcRn-binding affinities by BLI. Our measurements show that MFc4 binds FcRn with a *K*_D_ of 72.0 nM at pH 6.0 and 433.8 nM at pH 7.4 (Supplementary Figure 2). The discrepancy between the reported *K*_D_ of 5 nM and our measured *K*_D_ of 72.0 nM at pH 6.0 may be due to differences in experimental conditions, although the *K*_D_ values for another variant, MFc3, are similar between the two measurements (330 nM *vs*. 177 nM at pH 6.0) (Table 3). In our measurements, MFc4 showed a 6-fold difference in FcRn-binding affinity between pH 6.0 and pH 7.4. In contrast, three of our mFc variants (IgG1_mFc3, IgG1_mFc4, IgG1_mFc6) exhibited much greater pH responsiveness, with 213- to 445-fold differences in the affinity between the two pH values (Table 3). These mFc variants exhibit tight binding affinity at low pH (*K*_D_ of 31-88 nM), which is considerably stronger than the reported FcRn-binding avidity of dimeric IgG1 (*K*_D_ of 300 nM or 550 nM) [18, 23]. Such binding characteristics suggest that these mFc variants can effectively outcompete endogenous IgG1 and IgG4 molecules for FcRn binding in the endosome, while still being readily released from FcRn upon reaching the cell surface.

**Table 3.**
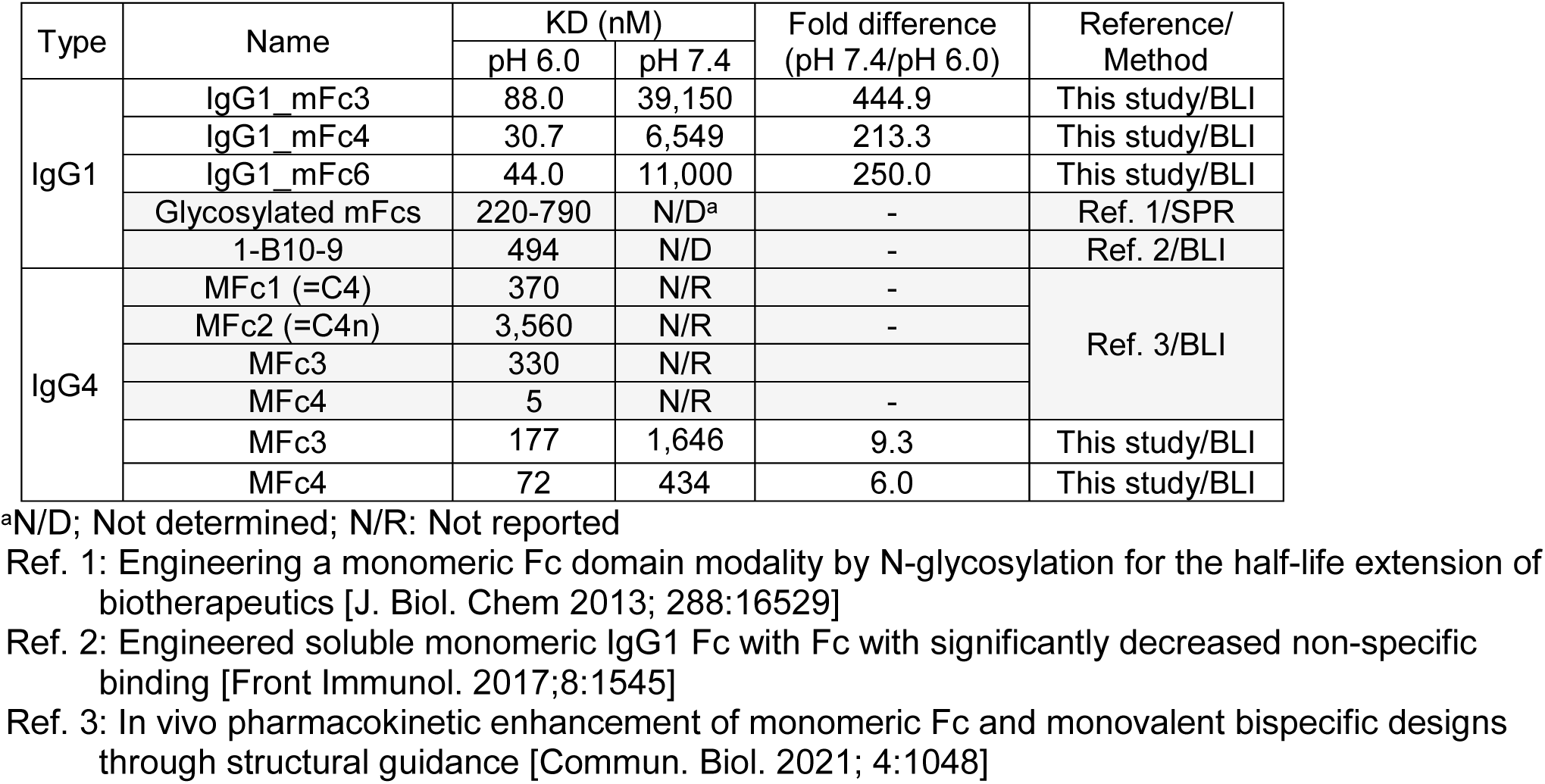
Comparison of human FcRn-binding affinities of mFc variants.

### Generation of IgG4 mFc and dimeric IgG Fc variants

Given the high sequence and structural similarity between the C_H3_ domains of IgG1 and IgG4, we introduced the same α2 sequence and Y407R mutation from IgG1_mFc2 into the human IgG4 Fc, generating IgG4_mFc2. This protein exhibited monomeric behavior (Supplementary Figure 3A) and remained soluble at concentrations of at least 25 mg/mL. Additionally, the amino acid substitutions present in IgG1_mFc3, which enhance FcRn-binding affinity in a pH-responsive manner, were also introduced into IgG4_mFc2, with the expectation of achieving similar binding properties. Indeed, the resulting variant, designated IgG4_mFc3, bound FcRn with a *K*_D_ of 110 nM at pH 6.0 and 24,510 nM at pH 7.4 (Supplementary Figure 3B). Therefore, transferring the amino acid substitutions present in other IgG1 mFc variants into IgG4_mFc2 would generate IgG4 mFc variants with FcRn-binding properties similar to those of the parent IgG1 mFc variants.

To assess whether the FcRn-affinity enhancing substitutions from IgG1_mFc3 could confer pH-responsive FcRn-binding avidity in a full-length antibody context, we introduced these substitutions into the Trastuzumab Fc. The resulting variant, designated IgG1_dFc3, exhibited a 7.6-fold increase in FcRn-binding avidity at pH 6.0 but only a 2.2-fold increase at pH 7.4, compared to the native dimeric Fc in Trastuzumab (Supplementary Figure 5). The fold increase in avidity at pH 6.0 is substantially lower than the 17-fold increase in affinity observed for the parental mFc variant, contrary to expectations. This discrepancy may be attributed to differences in conformational flexibility between the dimeric and monomeric Fc formats.

Nonetheless, the strong pH-selective FcRn-binding avidity of IgG1_dFc3 supports its potential as a dimeric Fc platform for extending the serum half-life of antibodies and other therapeutic modalities.

## Discussion

Engineered monomeric Fc molecules tend to aggregate at elevated concentration due to the exposure of the hydrophobic dimerization interface in the C_H3_ region [22, 31]. In this study, we demonstrated that just two substitutions, S354R and Y407R, can disrupt the Fc dimerization interaction but compromise solution behavior and protein stability. However, when combined with the additional remodeled helix α2, these modifications produced an mFc variant with significantly improved solution properties. This variant maintained a monomeric state under crystallization conditions, remained soluble at least at 25 mg/mL, and exhibited thermal stability with a melting temperature (T_m_) of 62.1 °C.

This IgG1_mFc2 variant was a starting scaffold to generate mFcs with enhanced pH-responsive FcRn-binding affinity through computational protein design and rational approach. A key implementation achieved by protein design method was the 4-residue extension of the loop composed of residues 284-286, which greatly enhanced the FcRn-binding at pH 6.0, likely by reinforcing the hydrophobic intermolecular interaction near the pH-responsive His310^Fc^-Glu115^FcRn^ pair. The robust pH-dependent FcRn-binding properties of IgG1_mFc3, IgG1_mFc4 and IgG1_mFc6, characterized by strong binding at acidic pH and rapid release at neutral pH, are essential for efficient Fc recycling and extended serum half-life *in vivo* [18, 24]. The three mFc variants exhibit two-digit nanomolar binding affinity at pH 6.0, but weak affinities at pH 7.4 (*K*_D_ ranging from 6,549 to 39,150 nM), which are well below the threshold (*K*_D_ of 860 nM) for dimeric Fc to exhibit an *in vivo* half-life comparable to that of native Fc [8], indicating their potential to extend the half-life of therapeutic modalities fused to these mFcs. It is noted that this threshold affinity is expected to be much lower of mFc (i.e., *K*_D_ << 860 nM) due to the absence of binding avidity for FcRn.

While dimeric Fc variants engineered for extended half-life have successfully been incorporated into therapeutic antibodies that are either clinically approved or in preclinical stages, such as antibodies with YTE or LS mutations [14, 32, 33], there have been no mFc variants developed to the clinical stage. The absence of clinically viable monomeric Fc-based therapeutics is largely attributed to multiple technical and biophysical challenges. These challenges include suboptimal pharmacokinetic properties, notably diminished serum half-life due to the loss of avidity for FcRn, decreased solubility and thermal stability associated with the exposure of hydrophobic residues upon monomerization, and lower yields during protein expression and purification processes [21, 31].

Together with high solubility, thermal stability, negligible immunogenicity, the robust pH-dependent FcRn-binding characteristics of a number of mFc variants developed in this study address the key barriers previously faced by monomeric Fc constructs. These properties strongly support the feasibility of these mFcs as viable platforms for therapeutic development, providing opportunities to develop mFc-fusions or antibody fragments with clinically relevant pharmacokinetics and developability profiles.

On the other hand, IgG1_mFc5 and IgG1_mFc9, identified through affinity tuning via variations in residues 433-434 or in the designed loop, exhibit potent FcRn-binding affinities at both pH values. Specifically, they show *K*_D_ values of 7.5 nM and 48.2 nM at pH 6.4, and 1,056 nM and 342 nM at pH 7.4, respectively (Table 1; Supplementary Figure 1). With these binding properties, the two mFc variants could be developed as therapeutic FcRn blockers for autoimmune diseases by tightly binding FcRn across both pH ranges, effectively inhibiting recycling of endogenous IgGs and thereby reducing serum levels of pathogenic autoantibodies

In conclusion, through protein design, we identified a novel and robust set of amino acid variations that convert native dimeric Fc into a stable monomeric state. In this mFc, we incorporated a designed loop that can serve as a versatile platform for generating a series of variants with diverse pH-dependent FcRn-binding profiles. Characterization of the generated mFcs indicates promising *in vivo* potential for extending the half-life of fused therapeutic modalities. Furthermore, the same substitutions can be applied to generate IgG4 mFc variants with strong pH-responsive FcRn-binding properties comparable to those of the parental IgG1_mFc. The same strategy can also be extended to produce dimeric IgG1 or IgG4 variants with enhanced pH-dependent FcRn-binding avidity, although further binding analyses are required.

## Methods

### Computational design

The reference sequences for wild-type IgG1 Fc and IgG4 Fc used in this study are based on UniProt entries P01857 (IgG1 allotype: G1m17,-1) and P01861 (IgG4), respectively. The structure of IgG1 Fc (PDB entry: 1L6X) was used as a template. Residues between 357 and 362 were inpainted with 7 to 10 amino acids (corresponding to one to four amino acid insertion). The output backbone structures were manually inspected, and a one-residue insertion was determined to be optimal. An Inpainting model that has the highest confidence score given by pLDDT (predicted local distance difference test) was selected, and the full coordinates of the inpainted segment were recovered by the relaxation protocol at the Cartesian coordinate space performed in the Rosetta software. The sequence of helix α2 was designed by the FastDesign protocol in Rosetta and by ProteinMPNN. Based on structure prediction of the designed models by AlphaFold2 and visual inspection, seven designs were selected for protein production and biochemical validation.

### Production of mFc variants

DNA fragments encoding each mFc were synthesized (IDT) and cloned into a vector derived from the pTT5 vector (Addgene) which contains the C_H2_-C_H3_ sequence of IgG. The resulting vectors were introduced into CHO-S cells (Gibco) at a density of 6×10^6^ cells/mL using ExpiFectamine (Gibco) to express mFc proteins. The cells were grown in the ExpiCHO expression medium (Gibco) for 10 days. The culture supernatant was collected by centrifugation at 12,000g for 1 h at 4 °C, diluted by half with Protein A binding buffer (PBS, pH 7.4), loaded onto an open column containing Protein A resin (Genscript), and eluted with Protein A elution buffer (0.1 M glycine, pH 3.0). The eluent was immediately neutralized with neutralizing buffer (1M Tris-HCl, pH 8.5), and further purified using a HiLoad 26/60 Superdex 75 column (Cytiva) equilibrated with PBS. Protein concentration was measured using a NanoDrop™ One spectrophotometer (Thermo Scientific) after concentrating the sample to approximately 1 mg/mL.

### Production of Trastuzumab Fab-mFc, hGH-mFc and hGH

DNA fragments encoding the V_H_-C_H1_ or V_L_-C_L_ regions of Trastuzumab were synthesized (IDT) and cloned into a pTT5-derived vector containing the mFc-coding sequence (pTT5-mFc vector) or the unmodified pTT5 vector, respectively. A DNA fragment encoding hGH (residues 27–217) was synthesized (IDT) and cloned into either the pTT5-mFc vector or the pTT5 vector with a C-terminal 6×His tag. The resulting vectors were used to produce the proteins from CHO-S cells in 25 mL of the ExpiCHO expression medium. Expression and protein purification of Trastuzumab Fab-mFc and hGH-mFc followed a protocol established for the mFc variant production. Purification of hGH was essentially identical, except that HisPur Ni-NTA resin (Thermo Scientific) was used instead of MabSelect™ PrismA resin.

### Crystallization, structure determination and refinement

Optimized crystallization conditions for each IgG1_mFc protein were established by hanging-drop vapor diffusion at 20 °C, using 2 μL drops prepared by mixing equal volumes of protein and reservoir solutions. All crystals were flash-frozen in liquid nitrogen after cryoprotection.

The crystallization and cryoprotection conditions are as follows;

IgG1_mFc1: Protein (5 mg/mL); reservoir solution: 1.7 M ammonium sulfate, 100 mM HEPES (pH 7.5); cryoprotection: reservoir solution + 10% (v/v) glycerol.
IgG1_mFc3: Protein (4.5 mg/mL); reservoir solution: 200 mM sodium acetate, 100 mM Tris (pH 8.5), 25% (v/v) PEG 4000; cryoprotection: reservoir solution + 5% (v/v) ethylene glycol.
IgG1_mFc6: Protein (4.5 mg/mL); reservoir solution: 200 mM sodium acetate, 100 mM Tris (pH 8.5), 30% (v/v) PEG 4000; cryoprotection: reservoir solution + 2.5% (v/v) ethylene glycol.
IgG1_mFc9: Protein (3.4 mg/mL); reservoir solution: 1.2 M sodium/potassium phosphate (pH 8.2); cryoprotection: reservoir solution + 10% (v/v) ethylene glycol.

X-ray diffraction data were collected at beamline 5C of Pohang Accelerator Laboratory (Republic of Korea) and processed with HKL2000 [34]. Structures were determined by molecular replacement, and refined using COOT [35] and PHENIX [36] (Supplementary Table 1).

### Biolayer interferometry

For BLI measurements at pH 6.0, mFc variants were concentrated and buffer-exchanged into 150 mM NaCl, 50 mM MES buffer (pH 6.0) using an Amicon centrifugal filter (Merck Millipore). BLI experiments were performed using an Octet R8 system (Sartorius). Biotinylated FcRn (ACRObiosystems) at 30 nM (1.5 ng/µL), prepared in either PBS buffer (pH 7.4) or 150 mM NaCl, 50 mM MES buffer (pH 6.0), was loaded onto streptavidin biosensor tips (Sartorius). The loaded sensors were then equilibrated in Kinetics Buffer (Sartorius) for 150 seconds. To measure the *K*_D_, each mFc variant at seven different concentrations was subjected to BLI runs with an association step (for 360 s) followed by a dissociation step (for 360 s). Binding kinetics were analyzed using the Octet BLI Analysis 12.2.2.4 software package (Sartorius).

### Circular dichroism spectroscopy

Purified proteins were diluted to 0.5 mg/mL in phosphate-buffered saline (PBS, pH 7.4) and placed into a 1 mm path-length quartz cuvette. Temperature-dependent unfolding was monitored using a JASCO J-1500 circular dichroism (CD) spectropolarimeter (JASCO), gradually increasing the temperature from 40 °C to 90 °C. Ellipticity values at 206 nm were recorded at 2 °C increments and converted to mean residue ellipticity (MRE). Thermal denaturation curves were analyzed by fitting MRE data to a Boltzmann sigmoidal equation using GraphPad Prism 6 software.

### Immunogenicity assessment

Peripheral blood samples were collected from healthy donors after obtaining informed consent under a protocol approved by the Institutional Review Board of Samsung Medical Center (IRB No. 2010-04-039-077). PBMCs were isolated by density centrifugation using Ficoll-Plaque Plus (Cytiva), and red blood cells were lysed with RBC lysis buffer (iNtRON Biotechnology). Primary human PBMCs were cultured in RPMI-1640 medium supplemented with 5% human AB serum, 25 mM HEPES, 2 mM L-glutamine, 100 units/ml penicillin and 100 μg/ml streptomycin. To assess immunogenicity, PBMCs were seeded in a flat-bottom culture plate at a concentration of 1 x 10^6^ cells/mL and challenged with mFcs. After 24 h of stimulation, culture supernatants were collected and analyzed by ELISA for quantification of pro-inflammatory cytokines using the IFN-γ Quantikine kit (R&D Systems, cat. DIF-50C), IL-2 Quantikine kit (R&D Systems, cat. D2050), and TNF-α Quantikine kit (R&D Systems, cat. DTA00D), according to the manufacturer’s instructions.

## Data availability

The coordinates of the four mFc structures will be deposited in the Protein Data Bank with the conditions of immediate release upon publication.

## Acknowledgement

This study utilized the Beamline 5C at the Pohang Accelerator Laboratory, Republic of Korea. The research was supported by the National Research Foundation of Korea (Grant Numbers. RS-2023-NR077287, RS-2024-00440039, RS-2024-00467046 and RS-2024-00508861) and by the KAIST Convergence Research Institute Operation Program.

## Author contributions

B.-H.O. and B.J. designed and directed the work. B.-H.O. and S.-B.I performed the computational designs. J.-Y.J., H.L., S.C., S.S., Y.-G.K. performed the protein production, crystallographic and other biochemical work. M.-J. A. directed and B.M.K. performed immunogenicity assay. B.-H.O., B.S.J., J.-Y.J. and S.C. wrote the original draft. All authors reviewed and accepted the manuscript.

