## Supplementary tables and figures for "Computational design of monomeric Fc variants with distinct pH-responsive FcRn-binding profiles"

**Supplementary Table 1. X-ray data collection and structure refinement statistics**

| Data Collection | **IgG1_mFc1** | **IgG1_mFc3** | **IgG1_mFc6** | **IgG1_mFc9** |
|---|---|---|---|---|
| Space group | $P\ 2_1\ 2_1\ 2_1$ | $P\ 2_1$ | $P\ 2_1$ | $C222_1$ |
| Unit cell dimensions | | | | |
| a, b, c (Å) | 72.6, 76.1, 110.0 | 71.0, 58.0, 71.4 | 70.8, 57.7, 71.1 | 89.2, 105.1, 71.4 |
| α, β, γ (°) | 90.0, 90.0, 90.0 | 90.0, 111.6, 90.0 | 90.0, 114.6, 90.0 | 90.0, 90.0, 90.0 |
| Wavelength (Å) | 1.0000 | 1.0000 | 1.0000 | 1.0000 |
| Resolution (Å) | 38.06 - 2.14 (2.22 - 2.14)[a] | 29.46 – 1.60 (1.66 - 1.60) | 28.87-1.77 (1.83 - 1.77) | 28.61-1.75 (1.80 - 1.70) |
| R-merge (%) | 0.08 (1.07) | 0.14 (0.97) | 0.08 (14.4) | 0.06 (0.59) |
| I/σ(I) | 24 (1.9) | 10.5 (3) | 15.4 (1.6) | 26.8 (5.1) |
| Completeness (%, > 0 σ) | 99.8 (98.6) | 99.5 (99.1) | 95.4 (89.5) | 90.81 (98.8) |
| Redundancy | 12.8 | 6.8 | 6.7 | 13.1 |
| Refinement | | | | |
| No. of reflections | 31644 (2683) | 71015 (7046) | 48552 (4399) | 31,079 (3,354) |
| $R_{work}$ / $R_{free}$ (%) | 22.7 / 27.6 | 19.6 / 22.8 | 21.4 / 26.0 | 22.6 / 25.9 |
| R.m.s deviations | | | | |
| bond (Å) / angle (°) | 0.010/1.18 | 0.007 / 0.89 | 0.008 / 0.93 | 0.007 / 0.87 |
| Average B-values (Å$^2$) | 81.0 | 19.0 | 32.0 | 26.36 |
| Ramachandran plot (%) | | | | |
| Favored/Additionally allowed | 97.6 / 2.16 | 98.6 / 1.4 | 99.1 / 0.9 | 100.0 / 0.0 |
| Outliers | 0.24 | 0.0 | 0.0 | 0.0 |

[a]The numbers in parentheses are the statistics from the highest resolution shell.

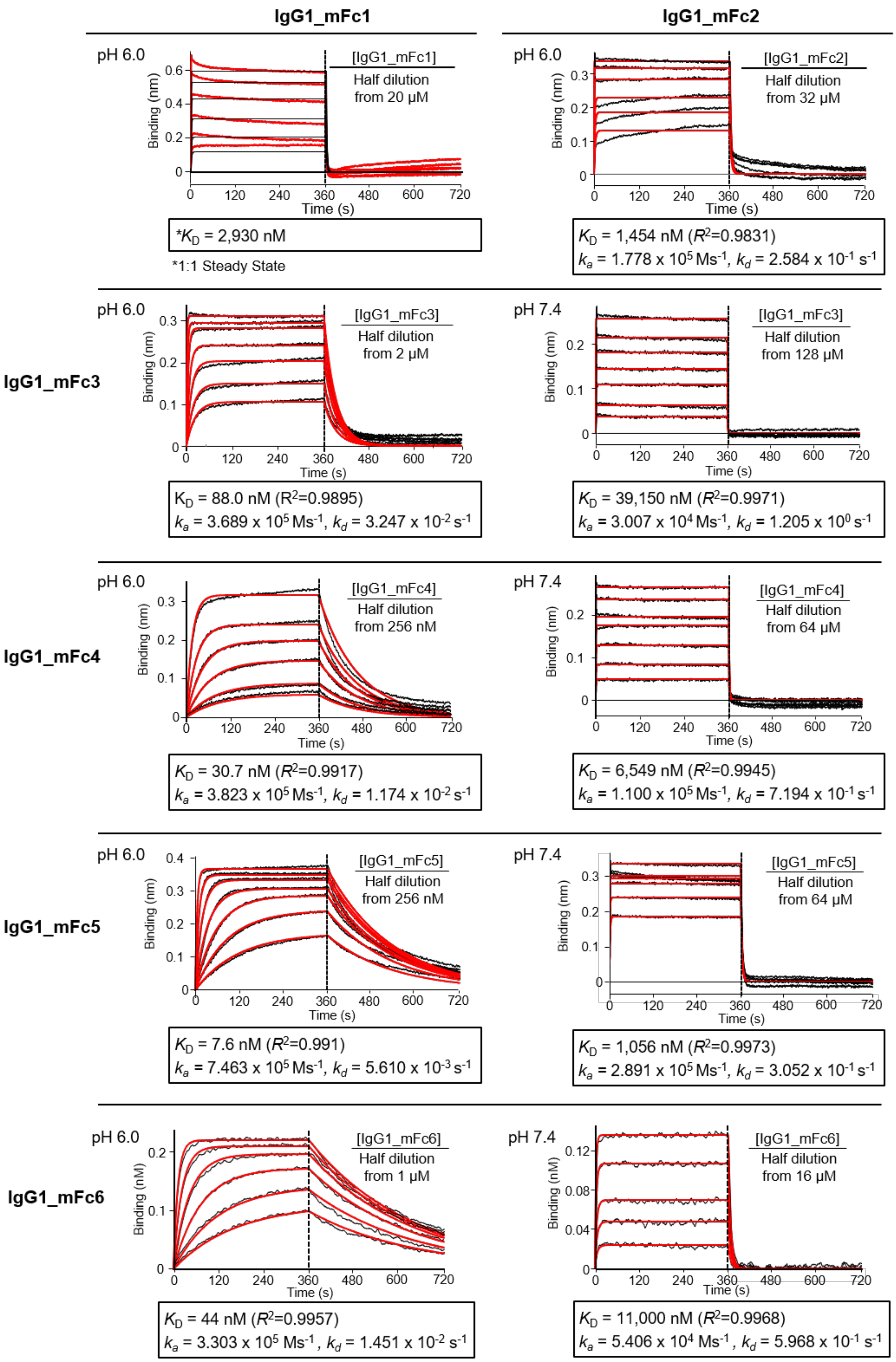
IgG1_mFc1
IgG1_mFc2
pH 6.0
[IgG1_mFc1] Half dilution from 20 µM
$*K_D$ = 2,930 nM
*1:1 Steady State
pH 6.0
[IgG1_mFc2] Half dilution from 32 µM
$K_D$ = 1,454 nM ($R^2$=0.9831)
$k_a$ = 1.778 x $10^5$ $Ms^{-1}$, $k_d$ = 2.584 x $10^{-1}$ $s^{-1}$
IgG1_mFc3
pH 6.0
[IgG1_mFc3] Half dilution from 2 µM
$K_D$ = 88.0 nM ($R^2$=0.9895)
$k_a$ = 3.689 x $10^5$ $Ms^{-1}$, $k_d$ = 3.247 x $10^{-2}$ $s^{-1}$
pH 7.4
[IgG1_mFc3] Half dilution from 128 µM
$K_D$ = 39,150 nM ($R^2$=0.9971)
$k_a$ = 3.007 x $10^4$ $Ms^{-1}$, $k_d$ = 1.205 x $10^0$ $s^{-1}$
IgG1_mFc4
pH 6.0
[IgG1_mFc4] Half dilution from 256 nM
$K_D$ = 30.7 nM ($R^2$=0.9917)
$k_a$ = 3.823 x $10^5$ $Ms^{-1}$, $k_d$ = 1.174 x $10^{-2}$ $s^{-1}$
pH 7.4
[IgG1_mFc4] Half dilution from 64 µM
$K_D$ = 6,549 nM ($R^2$=0.9945)
$k_a$ = 1.100 x $10^5$ $Ms^{-1}$, $k_d$ = 7.194 x $10^{-1}$ $s^{-1}$
IgG1_mFc5
pH 6.0
[IgG1_mFc5] Half dilution from 256 nM
$K_D$ = 7.6 nM ($R^2$=0.991)
$k_a$ = 7.463 x $10^5$ $Ms^{-1}$, $k_d$ = 5.610 x $10^{-3}$ $s^{-1}$
pH 7.4
[IgG1_mFc5] Half dilution from 64 µM
$K_D$ = 1,056 nM ($R^2$=0.9973)
$k_a$ = 2.891 x $10^5$ $Ms^{-1}$, $k_d$ = 3.052 x $10^{-1}$ $s^{-1}$
IgG1_mFc6
pH 6.0
[IgG1_mFc6] Half dilution from 1 µM
$K_D$ = 44 nM ($R^2$=0.9957)
$k_a$ = 3.303 x $10^5$ $Ms^{-1}$, $k_d$ = 1.451 x $10^{-2}$ $s^{-1}$
pH 7.4
[IgG1_mFc6] Half dilution from 16 µM
$K_D$ = 11,000 nM ($R^2$=0.9968)
$k_a$ = 5.406 x $10^4$ $Ms^{-1}$, $k_d$ = 5.968 x $10^{-1}$ $s^{-1}$
Binding (nm)
Binding (nM)
Time (s)
0
120
240
360
480
600
720

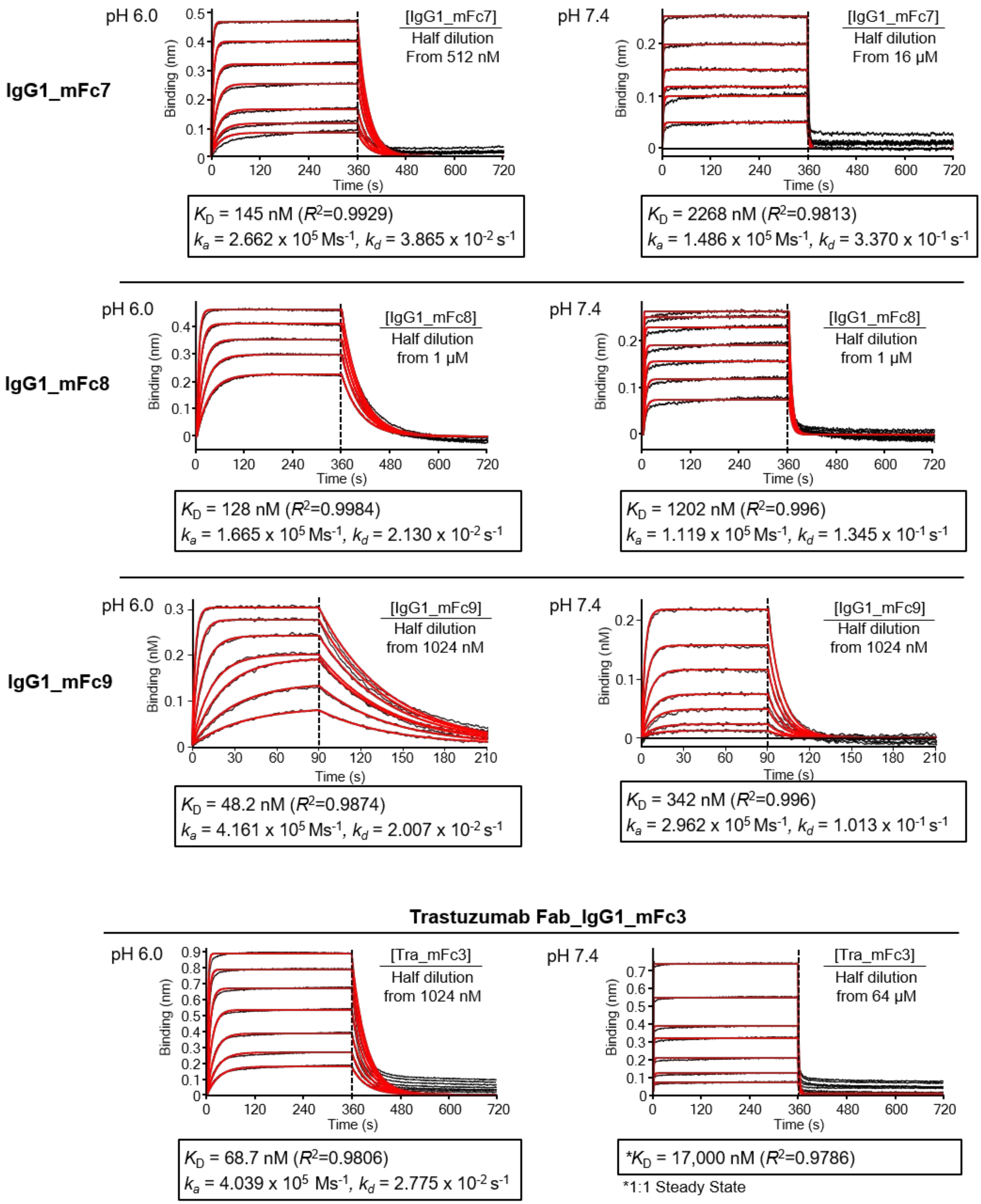


**Supplementary Figure 1. BLI runs for all 9 mFc variants with $k_a$ and $k_d$ values and IgG1_mFc3 fused to Trastuzumab Fab.**

FcRn-binding affinity of the indicated mFc variants were measured at pH 6.0 and pH 7.4. The deduced $K_D$ values are shown together with $k_a$ and $k_d$ values. Fitted curves are displayed in red. For IgG1_mFc1 at pH 6.0 and Trastuzumab Fab_IgG1_mFc3 at pH 7.4, curve fitting was performed using a 1:1 steady state model.

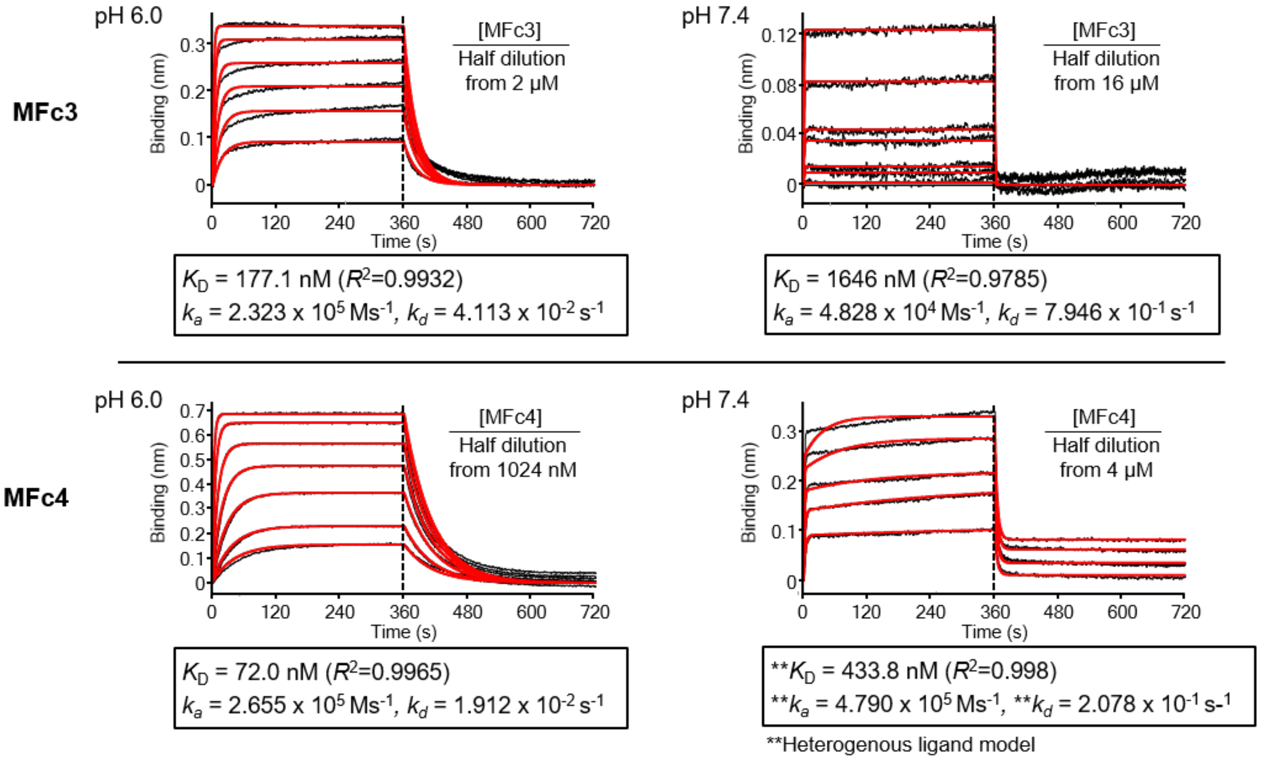


**Supplementary Figure 2. BLI runs for MFc3 and MFc4.**
FcRn-binding affinity of the indicated mFc variants were measured at pH 6.0 and pH 7.4. The deduced $K_D$ values are shown together with $k_a$ and $k_d$ values. Fitted curves are displayed in red. For MFc4 at pH 7.4, curve fitting was performed using a heterogenous ligand model.

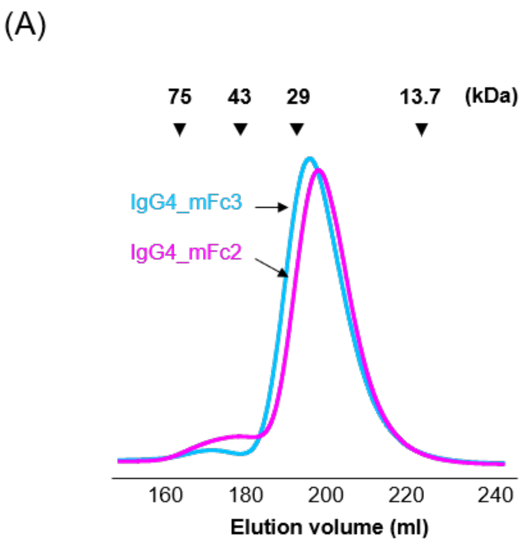


| Name | Dimer size | Monomer |
| --- | --- | --- |
| IgG4_mFc2 | 9.2 % | 90.8 % |
| IgG4_mFc3 | 3.9 % | 96.1 % |

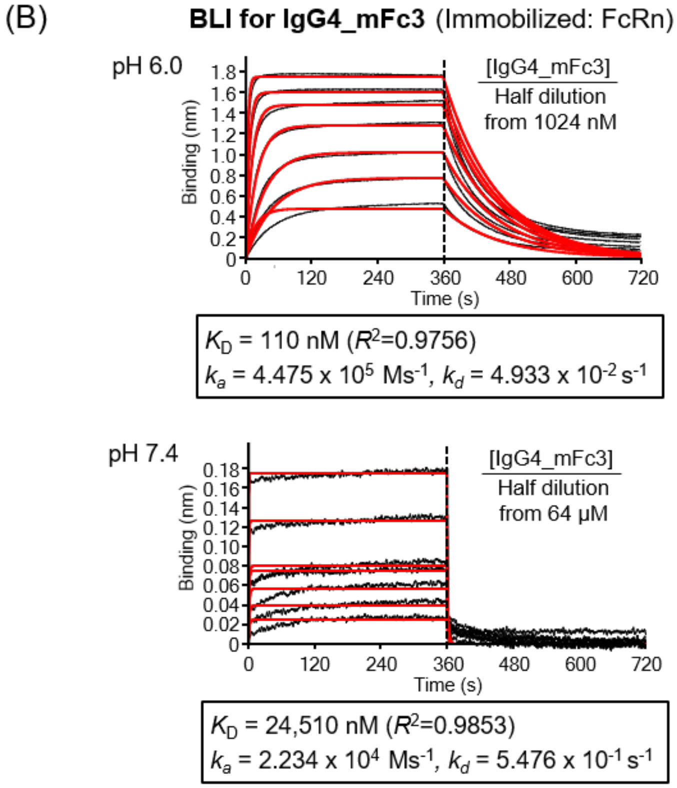


**Supplementary Figure 3. Solution behavior of IgG4 mFc variants and binding affinity quantification.**

(A) Elution profile. IgG4_mFc2 and IgG3_mFc3, after elution from protein A resin, were concentrated to 10 mg/mL and loaded onto a Sephadex 75 column. Size marker proteins are indicated by triangles.

(B) BLI runs for IgG4_mFc3. FcRn-binding affinities were measured at pH 6.0 and pH 7.4. The deduced $K_D$ values are shown together with $k_a$ and $k_d$ values. Fitted curves are displayed in red.

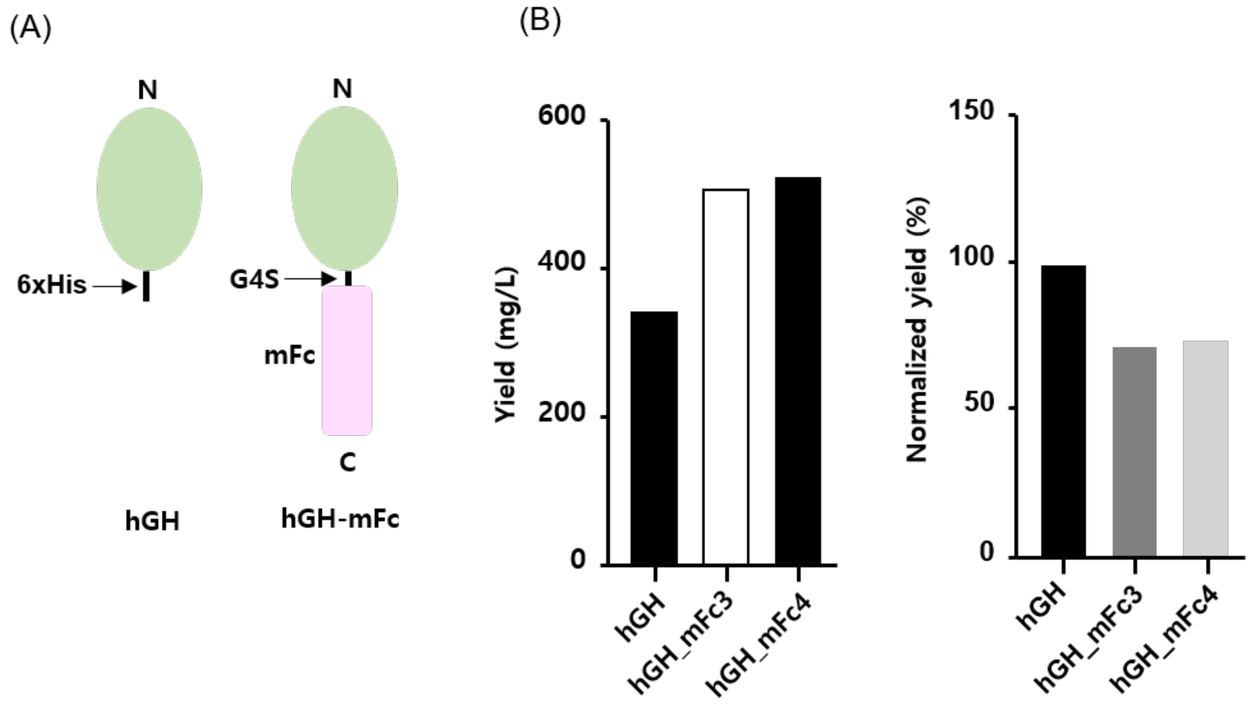


**Supplementary Figure 4. Comparison of hGH and hGH-mFc fusion protein production.**
(A) Constructs. hGH was fused to the N-terminus of mFc via a GGGGS (G4S) linker.
(B) Protein yield. The indicated proteins were expressed in 25 mL CHO-S cultures and purified. Shown are the total yields and yields normalized to molecular weight.

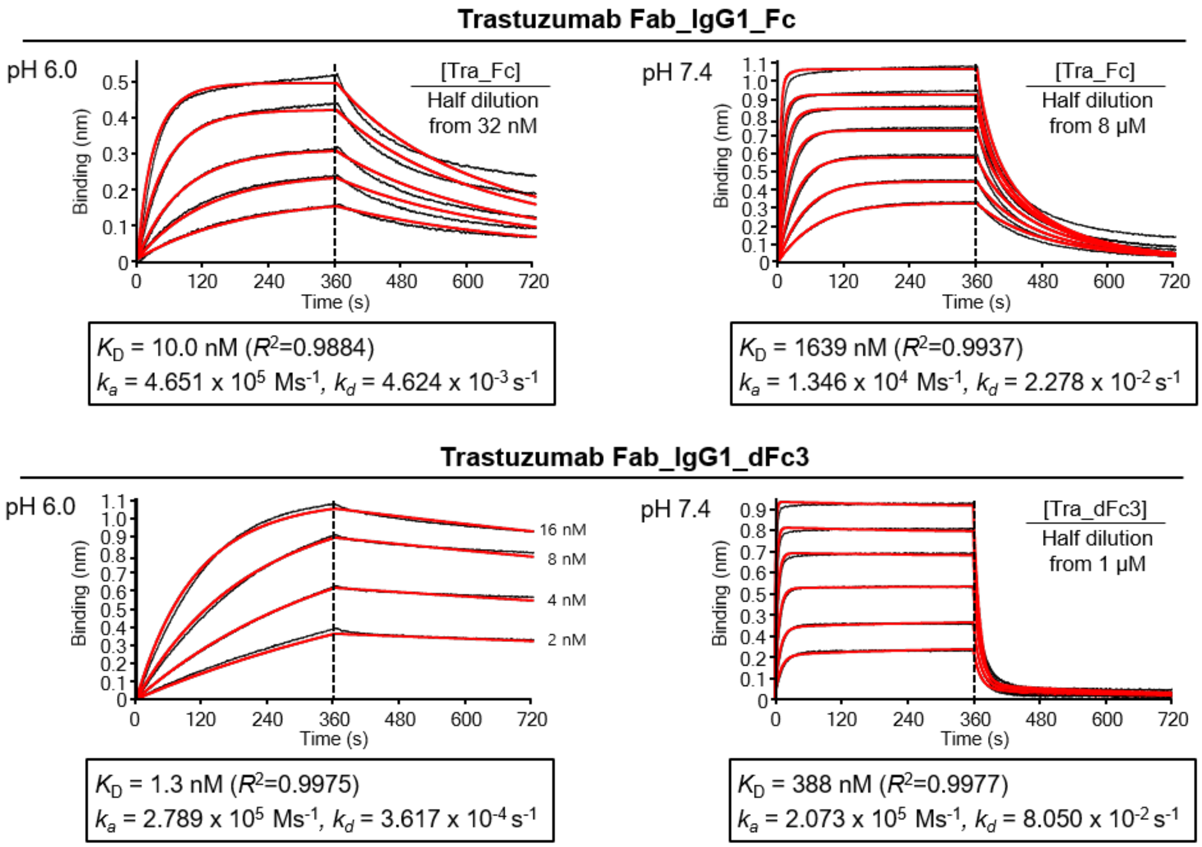


| Parental mFc | Dimeric Fc | $K_D$, nM (Avidity) | | Fold difference (pH 7.4/pH 6.0) |
|---|---|---|---|---|
| | | pH 6.0 | pH 7.4 | |
| - | IgG1 Fc | 10.0 | 1639 | 164 |
| IgG1_mFc3 | IgG1_dFc3 | 1.3 | 388 | 298 |

**Supplementary Figure 5. Comparison of the binding avidities of native dimeric Fc and dimeric Fc variant in the Trastuzumab background.**
FcRn-binding avidities were measured at pH 6.0 and pH 7.4. The deduced $K_D$ values are shown together with $k_a$ and $k_d$ values and compared in a table format. Curve fitting was performed using a bivalent analyte model.
